# A large cohort study (n = 591) on the impact of the presence or absence of the interthalamic adhesion: cognitive, neuroimaging, and genetic results

**DOI:** 10.1101/2024.09.20.614108

**Authors:** Julie P. Vidal, Alexa Gouarderes, Marie Stéphanie Rabenantenaina, Patrice Péran, Jérémie Pariente, Lola Danet, Emmanuel J. Barbeau

## Abstract

Both thalami can be connected by an Interthalamic Adhesion (IA). The extent of its presence varies among individuals and remains poorly understood. This study examines the IA’s prevalence, anatomical variations, genetic determinants, and cognitive associations. Data from 591 healthy subjects (25-35 years) from the Human Connectome Project were analyzed and grouped as monozygotic (MZ) twins, dizygotic (DZ) twins, non-twin siblings, and unrelated individuals. MRI was used to characterize the IA, while neuropsychological assessments and Freesurfer parcellations were used to assess cognition and anatomical differences between subjects with or without an IA. The IA was absent in 12.7% of subjects, more commonly in males (20.0%) than females (6.3%), with no significant differences in age, education, or cognition between those with and without an IA. IA absence was associated with increased cerebrospinal fluid volumes, enlarged third ventricles, and thinning in several cortical areas. Genetic analysis revealed a higher concordance of IA presence among MZ twins (96%) than in other groups, indicating a strong genetic influence. The remaining 4% discrepancy was observed in male pairs only. This study underscores the genetic basis of IA, highlighting sexual dimorphism and neuroanatomical differences associated with its absence, though it does not affect cognition in healthy individuals.

## 1. Introduction

The thalamus is a bilateral nuclear complex located in the diencephalon (Herrero et al. 2002). It plays a pivotal role in processing and transmitting sensory information to the cerebral cortex. It also contributes to higher-order functions such as attention, consciousness, and memory through cortico-cortical connections (Guillery et al. 2002). An interthalamic adhesion (IA), located on the median border across the third ventricle, can connect both thalami (Davie and Baldwin 1967; Malobabic et al. 1987; Pavlovic et al. 2020; Patra et al. 2022).

IA studies are scarce, and much remains to be investigated to better understand its role. In addition, this structure is absent in approximately 20% of healthy subjects, more often in males (17%) compared to females (9%) (Rabl 1959; Malobabic et al. 1987; Nopoulos et al. 2001; ErbaLcı et al. 2002; Takahashi et al. 2008; Pavlovic et al. 2020; Borghei et al. 2020, 2021; Wong et al. 2021; Sahin et al. 2023; Vidal et al. 2024). This adds to the mystery of this structure as it is rare for a brain structure to be sometimes present and sometimes absent in the healthy population. Commonly presented as a single adhesion in the anterosuperior quadrant (Davie and Baldwin 1967; Malobabic et al. 1987; Pavlovic et al. 2020; Patra et al. 2022), the IA can also vary in shape. Anatomical variants include a broader form found in 18% of cases, more frequently in females, duplications in 2-10% of instances, and rare forms like bilobar, multiple, tubular, or rudimentary IA seen in less than 3% of subjects (Viller et al. 1887; Malobabic et al. 1987; Herrero et al. 2002; Guillery et al. 2002; Miller et al. 2008; Tresniak et al. 2011; Sahin et al. 2023; Tsutsumi et al. 2021). In around 2 to 3% of cases, the “kissing thalami” phenomenon, where both thalami adhere closely, can sometimes obscure the characterization of the IA on usual MRI (Miller et al. 2008; Borghei et al. 2020, 2021; Vidal et al. 2024).

The IA is a white matter tract. The precise connectivity supported by the IA remains unclear. However, diffusion imaging and tractography studies suggest an association with the anterior thalamic radiations, projecting towards the orbitofrontal cortex and medial frontal region (Damle et al. 2017; Kochanski et al. 2018; Sahin et al. 2023). In the absence of IA, fibers might cross via the posterior commissure, suggesting compensatory pathways (Kochanski et al. 2018). The IA is also connected to regions like the amygdala, hippocampus, entorhinal cortex, insula, pericalcarine cortex, cuneus, nucleus accumbens, caudate nucleus, lateral habenula, and posterior commissure (Borghei et al. 2021; Sahin et al. 2023). These connections indicate that the IA could play a significant role in interhemispheric communication.

The functional role of the IA is debated, with some researchers viewing it as a vestigial structure (Viller et al. 2018). The inconsistency of its presence among individuals has contributed to this perception. However, emerging research suggests potential cognitive implications, particularly in areas such as attention (Damle et al. 2017; Borghei et al. 2020), executive functions (Trzesniak et al. 2016; Borghei et al. 2020), and verbal memory (Trzesniak et al. 2016). Recent studies indicate that while healthy individuals show no significant cognitive differences related to IA presence, patients with thalamic strokes exhibit fewer impairments in verbal memory and executive functions if they possess an IA. This suggests a protective cognitive role for the IA, possibly aiding in functional reorganization and compensatory mechanisms post-stroke (Vidal et al. 2024). Despite these findings, the role of the IA necessitates further research.

Debates continue about the reasons behind the absence of IA, including potential correlations with age. A 3D transvaginal neurosonography study on fetuses demonstrated an IA presence of 100%, and in an MRI study on teenagers, this prevalence was maintained (Takahashi et al. 2009a; Birnbaum et al. 2018). Accordingly, a progressive thinning and elongation of IA in an oldest cohort of patients has been noted in an MRI study (Sen et al. 2005) and broad IA are found to be more frequent in subjects less than 40 years old (Trzesniak et al. 2016; Tsutsumi et al. 2021). However, no relationship between mean patient age and either prevalence or the length of the IA was observed in a meta-analysis (Trzesniak et al. 2011).

Different studies have suggested a relationship between IA absence, or shorter length, and ventricle enlargement (Davie and Baldwin 1967; Samra et al. 1968; Snyder et al. 1998; Meisenzahl et al. 2002; Takahashi et al. 2010; Borghei et al. 2021). Subjects without an IA exhibit significantly larger third ventricles compared to those with an IA in both schizophrenia and healthy individuals. Additionally, a negative correlation exists between third ventricle volume and IA length in controls and bipolar disorder patients (Takahashi et al. 2009a, 2010). As the IA develops alongside the ventricular system (Rosales and al. 1968; O’Rahilly and Muller 1990; Snyder et al. 1998), these findings suggest that IA anomalies and ventricle enlargement could indicate abnormal midline structure development. This is supported by the frequent occurrence of IA abnormalities with other midline malformations such as Chiari II malformation (Miller and al. 2008; Etus and al. 2017), hydrocephalus (Yamasaki and al. 1995; Kanemura and al. 2006), Cornelia de Lange syndrome (Whitehead and al. 2015), and diencephalic-mesencephalic junction dysplasia (Severino and al. 2017). Among pediatric patients with midline abnormalities, IA absence is four times higher compared to healthy groups (Whitehead and Najim 2020). Additionally, some studies suggest that increased CSF pressure in the third ventricle could compress and potentially rupture the IA, as illustrated by a case where the IA disappeared after hydrocephalus developed (El Damaty and al. 2017). It remains unknown whether neurological diseases associated with global atrophy, such as Alzheimer’s disease and its impact on the thalamus, are associated with IA shrinkage or reduced prevalence. Similarly, the effect of multiple sclerosis on the IA remains to be investigated.

Many studies have reported increased rates of IA absence or shorter length in neurodevelopmental and neuropsychiatric disorders such as schizophrenia, borderline personality disorder, major depression, and bipolar disorder [Andreasen and al. 1994; Snyder and al. 1998; Nopoulos et al. 2001; Takahashi et al. 2009a; 2009b; 2010. In addition, altered connections to the amygdala, frontal, and anterior cingulate cortex have been linked to these disorders. These structures have been demonstrated to be connected to the IA (Serap Monkul and al. 2005; Delbello and al. 2006; Chanen and al. 2008; Konarski and al. 2008). In summary, IA absence might be related to early neurodevelopmental abnormalities, particularly of midline structures, leading to abnormal neural networks, including the thalamus and related regions. Those abnormal neural networks could lead to functional and structural aberrations, such as deficits in the dopaminergic system. Indeed, recently, a meta-analysis revealed that IA absence is twice as likely in schizophrenia spectrum disorders. The authors suggested an IA role in the dopaminergic system (Trzesniak and al. 2011), in line with thalamic deficits observed in schizophrenia spectrum disorders etiopathology, as a low dopamine receptor binding or a volume reduction (Andreasen and al. 1994; Sánchez-González and al. 2005; Buchsbaum and al. 2006; Ettinger and al. 2013). Altogether, IA absence could indicate a higher risk of developing neuropsychiatric disorders (Nopoulos and al. 2001; Trzesniak and al. 2011).

Whether the presence or absence of IA is related to genetic factors remains unknown. Yet, genetic links are well-documented for neuropsychiatric disorders such as schizophrenia and bipolar disorders, with high heritability estimates (Lichtenstein and al. 2006; Zhang and al. 2006; Pantelis and al. 2014). A study on monozygotic and dizygotic twins estimated the heritability of bipolar disorder at 85% (McGuffin and al. 2003). Thus, a possibility could be that IA presence could be governed by genetic factors, also related to the spectrum of factors associated with neurodevelopmental and neuropsychiatric disorders.

In summary, many intriguing questions about the IA remain unresolved. We aim to advance our understanding of this brain structure by studying a large cohort of 591 healthy individuals, including monozygotic and dizygotic twins. Our primary hypothesis is that genetics play a significant role in influencing IA characteristics, as it is consistently linked to genetically related disorders. Additionally, we hypothesize that the absence of the IA should not lead to cognitive differences, given its absence in some individuals. To explore these hypotheses, magnetic resonance images from healthy young adults participating in the Human Connectome Project (HCP) were analyzed to: (1) identify the presence or absence of the IA, (2) classify its variants, and (3) examine associations with anatomical and neuropsychological differences. The subjects were categorized into four groups: monozygotic (MZ) twins, dizygotic (DZ) twins, non-twin siblings, and unrelated individuals.

## 2. Materials and Methods

### 2.1 Participants

Data were provided by the Human Connectome Project, WU-Minn Consortium (Principal Investigators: David Van Essen and Kamil Ugurbil; 1U54MH091657) funded by the 16 NIH Institutes and Centers that support the NIH Blueprint for Neuroscience Research; and by the McDonnell Center for Systems Neuroscience at Washington University (Van Essen and al. 2013). Participants were healthy individuals from families in Missouri, United States. Inclusion criteria required the absence of severe neurodevelopmental, neuropsychiatric, or neurological disorders, as well as diseases such as diabetes or hypertension. Twins born before 34 weeks of gestation and non-twins born before 37 weeks were excluded. Each subject underwent neuroimaging and neuropsychological examinations. Data from the HCP Young Adult sample, comprising 591 subjects selected from 1207 based on the availability of magnetic resonance imaging (MRI) data, including T1 and T2 sequences, were used. The sample included 306 females (51.8%) and 285 males (48.2%), with an average age of 28.5 years (SD = 3.6, range 22-35 years) and 15 years of education (SD = 1.8).

To investigate the genetic influences on the prevalence of the IA, subjects were divided into four groups: monozygotic twins (MZ), dizygotic twins (DZ), non-twin siblings, and unrelated individuals. The monozygotic and dizygotic relationships were genetically verified. For unrelated subjects, random pairings were created to form equivalent groups.

### 2.2 Neuropsychological Assessment

Relevant psychological and neuropsychological data were extracted from the HCP database. This included measures of depression, anxiety, and attention/hyperactivity from the Statistical Manual (DSM)-Oriented Scales (American Psychiatric Association and al. 2013). Cognitive abilities were assessed using the NIH Toolbox Cognition Battery (Weintraub and al. 2013), which included tests for working memory (List Sorting Working Memory Test), language (Oral Reading Recognition Test), and episodic memory (Picture Sequence Memory Test for non-verbal episodic memory, and the Penn Word Memory Test for verbal episodic memory). Data for depression, anxiety, and attention tests were missing for six individuals.

### 2.3 MRI Acquisition

MRI images were acquired using a Siemens 3T “Connectome Skyra” scanner. For T1 sequences, images were obtained with a 3T scanner using the following parameters: 3D MPRAGE sequences, voxel size = 0.7x0.7x0.7 mm, TE = 2.14 ms, TR = 2400 ms, TI = 1000 ms, flip angle = 8°, FOV = 224x224 mm, slice thickness = 0.7 mm isotropic. The parameters for the 3D T2-SPACE sequences were: voxel size = 0.7x0.7x0.7 mm, TE = 565 ms, TR = 3200 ms, FOV = 224x224 mm, slice thickness = 0.7 mm isotropic. Images were coregistered and reoriented to align the anterior and posterior commissures to standardize orientation and facilitate comparison.

### 2.4 IA and IA Variants Identification Protocol

Two independent raters characterized the presence, absence, and type of IA following a previously established protocol (Vidal and al. 2024). An extended video protocol to characterize the IA is publicly available [DOI: 10.13140/RG.2.2.35022.47689], and a training dataset can be sent on request. MRI images were reviewed using MRIcron software, starting with axial slices and then sagittal and coronal slices to refine the characterization of the IA. Confirmation was done using T2-weighted images. An IA was identified if a structure connecting both thalami was observed on at least one slice between the anterior and posterior commissures. Variants were classified as standard, broad, bilobar, or double based on specific criteria (Vidal and al. 2024). More specifically, a standard form is characterized by a thin adhesion comprising less than one-third of the thalamus on the current axial and coronal slices, with a small, rounded shape on sagittal slices. A broad variant is identified when comprising over one-third of the thalamus on the current axial and coronal slices. The bilobar form is distinguished on sagittal slices by a bilobed shape and on coronal slices two distinct adhesions can be visualized and must not be mistaken for a double variant. Finally, the double variant is identified by two independent IA, which needs to be confirmed on all slices. To improve the previous protocol and help better differentiate the standard form from a broad one, especially in the case of close thalami, it is possible to segment the IA on sagittal slices and identify the amount of shared matter between the two thalami.

Two raters (A.G. and M.S.R) conducted the identification process blindly to avoid biased sibling correlations. Cases with the “kissing thalami” phenomenon were excluded from statistical analyses. Consequently, the sibling or unrelated paired subject was also excluded from the genetic analyses, as our focus was on pairs. In cases of disagreement between the examiners, discussions were held to reach a consensus. A third expert opinion (J.P.V) was sought to resolve the differences if necessary. Inter-rater concordance was evaluated using Cohen’s Kappa coefficient for every 100 subjects.

### 2.5 Statistical Analyses

#### 2.5.1 Demographic data

The impact of handedness, biological sex, age, and education level on IA prevalence was examined using χ² tests for qualitative data and Student’s t-tests or Mann-Whitney tests for quantitative data. ANOVA or Kruskal-Wallis tests were used for variables with multiple categories.

#### 2.5.2 Prevalence of the IA, brain volumes and cortical thickness

FreeSurfer parcellations were used to extract volumes and cortical thicknesses for each subject, and their nomenclature was maintained for consistency. Given the effective differences between groups, Bayesian t-tests were conducted to identify anatomical differences between subjects with and without an IA. It can also give evidence for the absence of differences between groups and decrease bias from multiple comparisons. A Bayesian Factor (BF) < 0.3 indicates more evidence toward the lack of differences, while a BF > 3 moderate, BF> 10 strong, BF>30 very strong, and BF> 100 extreme indicates differences between each group. Volumes and thicknesses were corrected for intracranial volume multiplied by 10^6^. Cortical regions with a BF greater than 100 are represented for each hemisphere on a histogram. The 10 cortical regions by hemisphere with the highest BF are represented using MRIcroGL.

#### 2.5.3 Prevalence of the IA and neuropsychological consequences

An independent t-test was conducted to study the variation in neuropsychological test scores based on IA presence or absence.

#### 2.5.4 Prevalence of the IA and genetic causes

Logistic regressions were conducted to evaluate the association between the IA and genetics by comparing the four groups (MZ, DZ, non-twin, unrelated) and considering whether each paired subject possesses an IA or not. χ² tests with Bonferroni correction were used for pairwise comparisons. The same tests were performed for genetic analysis of IA variants, but only data related to the presence of IA in both paired subjects were retained. Finally, to anatomically and neuropsychologically compare siblings only differing by the presence or absence of the IA, the analyses were restricted to paired subjects when their adhesion status differed. In this aim, a paired Bayesian t-test was conducted to reduce bias due to low sample sizes, applied to volumes, cortical thickness, and psychological or cognitive test scores by group.

## 3. Results

### 3.1. Prevalence of the IA

The IA was characterized by two independent examiners with a mean concordance of 0.85 for the presence or absence and 0.79 for identifying anatomical variants using Cohen’s Kappa score. Out of the 591 subjects, 55 cases had kissing thalami, resulting in a final sample size of 536 subjects. Among these, 87.3% had an IA, while 12.7% did not. The absence of the IA was significantly higher in males (20%) compared to females (6.3%) (p < 0.001, χ² test = 22.6). No significant differences between subjects with and without an IA were observed for age, education, or manual laterality (Mann-Whitney U; p-value > 0.05).

### 3.2. Prevalence of IA variants

Recent developments in the study of the IA have included the analysis of anatomical variants (Tsutsumi and al. 2021). Following a previously established protocol (Vidal and al. 2024), we identified that among all subjects, 61% had a standard single IA, 25% had a broad variant, and less than 1% exhibited either a bilobar or double variant (Fig. 1).

**Figure 1:**
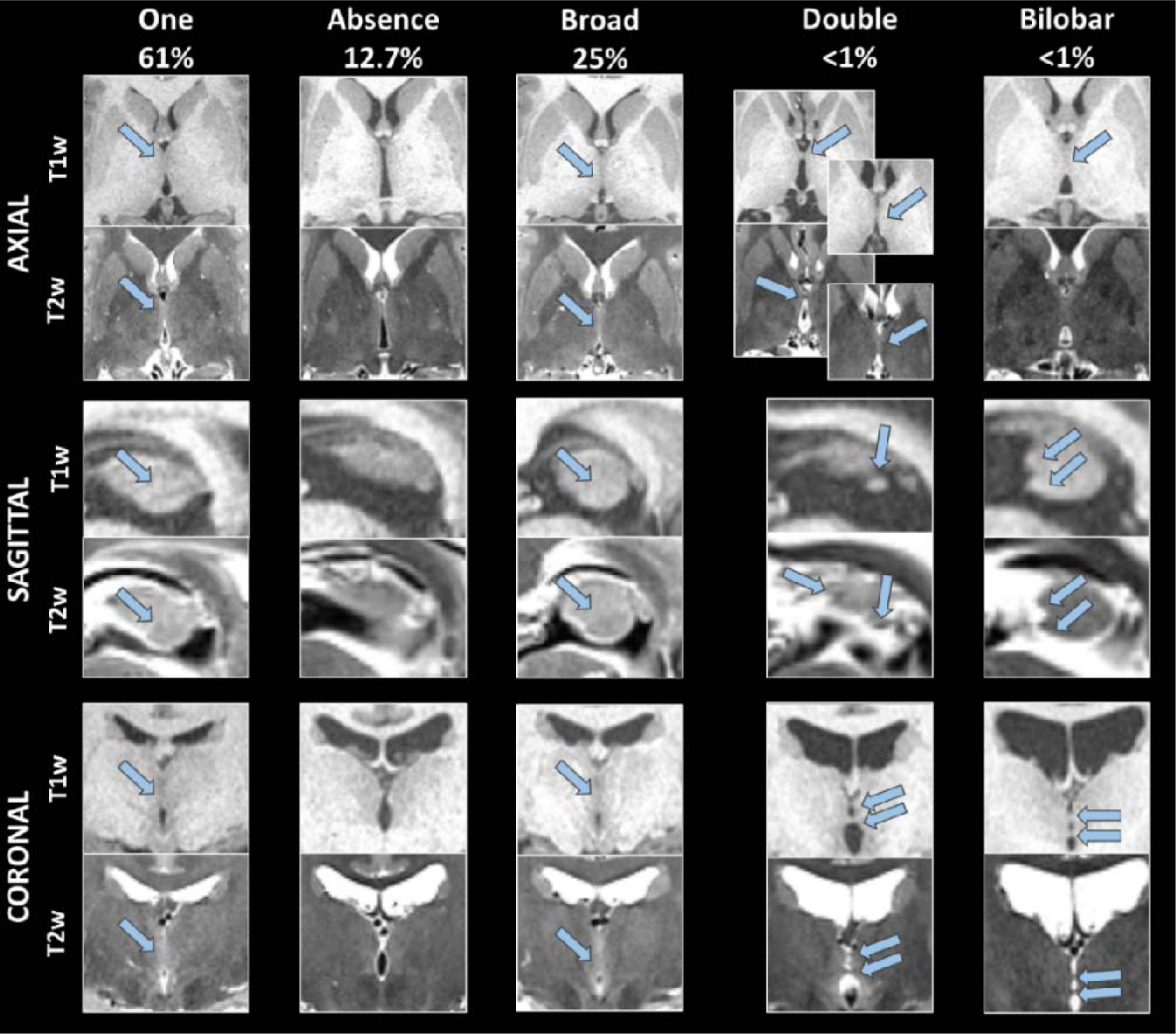
Typical single or absent IA and its most frequent anatomical variants (broad, double, bilobar) illustrated on the same slice of T1- or T2-weighted MRI. Blue arrows indicate the IA location. Note the lower contrast on T1w compared to T2w MRI, especially on sagittal slices, preventing the accurate location of the first double variant.

### 3.3. Prevalence of the IA and association with brain differences

Based on the literature about IA structural connectivity, we first address the hypothesis that anatomical differences exist between individuals with and without an IA. Using FreeSurfer parcellations, we sought to identify volume and cortical thickness variations associated with the IA presence. All comparisons (mean ± SD by groups and BF) are represented in Supplementary Table 1.

Subjects without an IA showed several differences compared to those with an IA (all differences are reported in detail in Supp. Tab. 1). Bayesian t-tests revealed moderate evidence for increased cerebrospinal fluid volumes (BF > 3) and extreme evidence for increased third ventricle volume (BF > 100) among subjects without an IA (Fig. 2A). This was associated with smaller corpus callosum (Fig. 2A) and bi-thalamic volume (Supp. Tab. 1). In addition, some brain regions showed cortical thinning, most notably the bilateral inferior and superior parietal, supramarginal, precentral and paracentral, superior frontal, precuneus, cuneus, and the banks of the superior temporal sulcus. While most differences were bilateral, some were unilateral. Figure 3 highlights all brain regions with BF10 > 100, and Figure 4 provides a schematic representation of these regions. Specifically focusing on the medial temporal lobe, differences were observed in the parahippocampal cortex bilaterally, right entorhinal cortex, and right temporal pole (Supp. Tab. 1).

**Figure 2:**
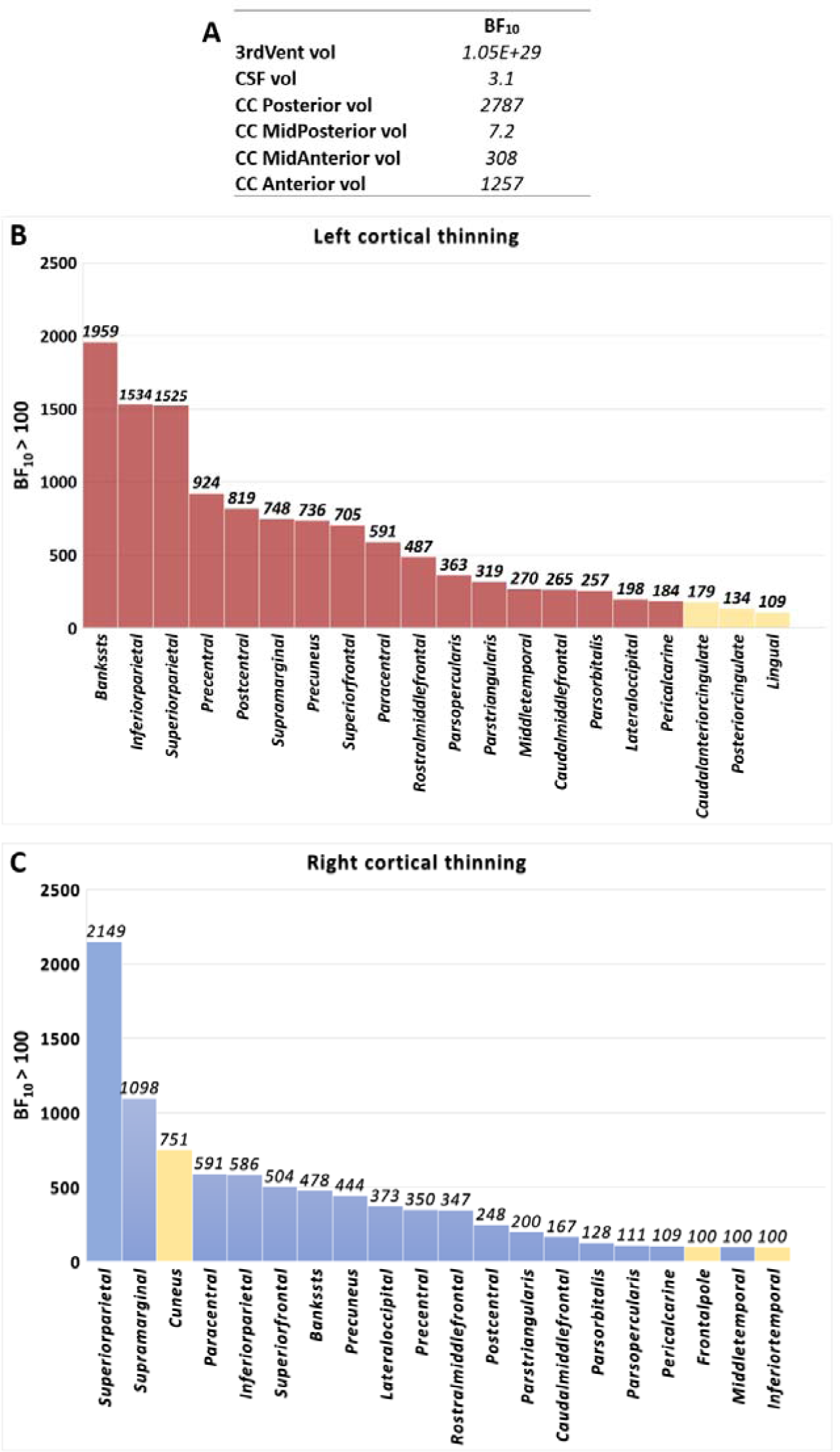
Comparisons of brain volumes and cortical thicknesses depending on the presence (N=468) or absence (N=68) of an IA, analyzed using a Bayesian t-test. The comparisons are presented for A) midline or general regions, B) left, or C) right brain regions. Only comparisons showing at least moderate evidence for a difference (BF > 3) are displayed. All volumes and thicknesses were corrected for intracranial volume and multiplied by a factor of 10^6^. Yellow bars represent regions with unilateral differences only. The names of the brain areas are from the nomenclature of FreeSurfer. CC: corpus callosum.

**Figure 3:**
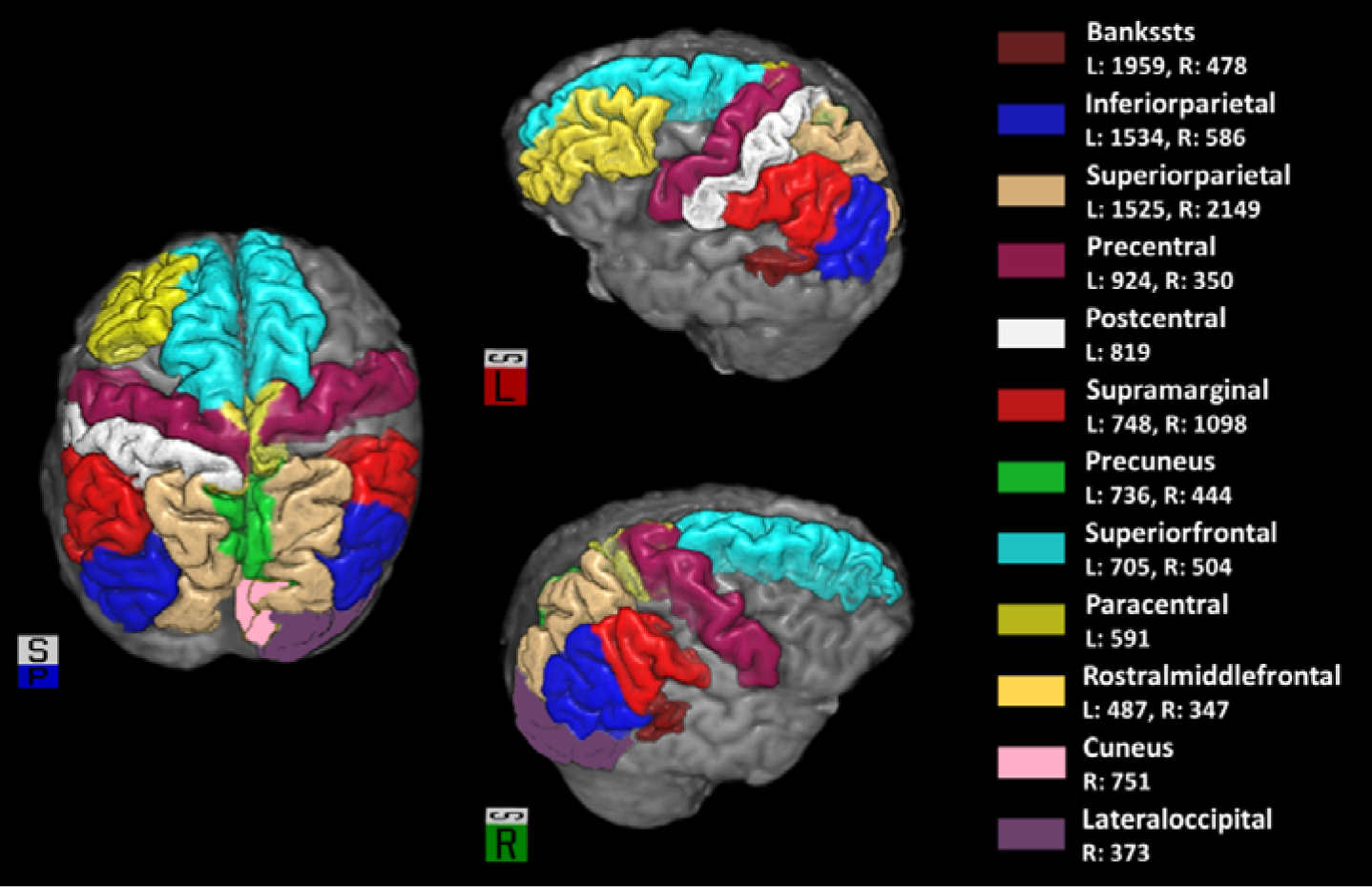
Illustration of the 10 cortical regions per hemisphere, as extracted from Freesurfer, that yielded the highest Bayes Factors (BF) in the previous analysis (Fig. 2). These regions demonstrate evidence of decreased cortical thickness in subjects without an IA compared to those with an IA. The figure provides a superior view (left), a left view (top), and a right view (bottom), with a cube indicating spatial orientation. The color legend identifies each region by name, along with the corresponding BF for each hemisphere.

**Figure 4:**
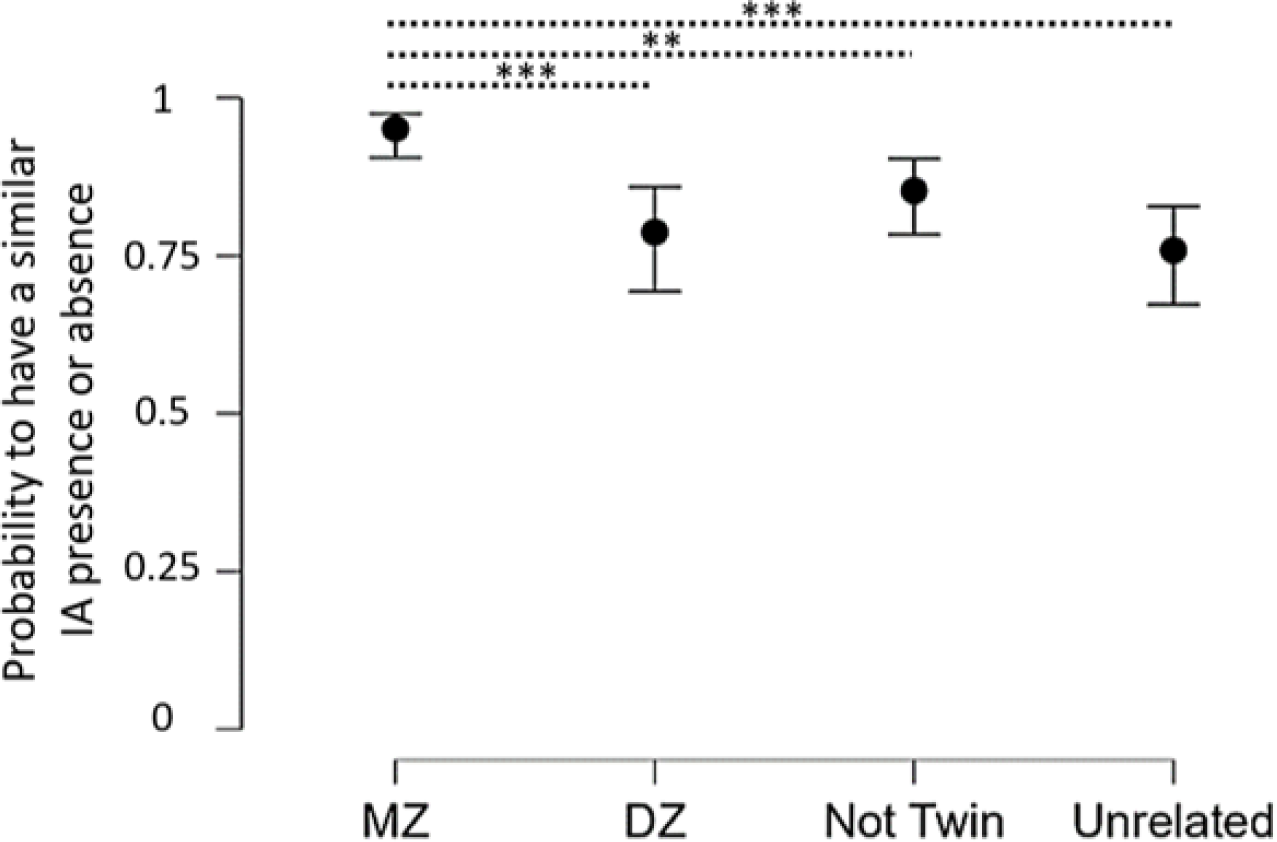
Probability to have the same IA between pairs and by genetic groups (MZ: monozygotic, DZ: dizygotic) ± 95% confidence interval. χ² p-values Bonferroni corrected: *** < 0.001, ** < 0.005.

### 3.4. Prevalence of the IA and association with affective and cognitive abilities

Given that the IA is absent in a large portion of the population and has been associated with brain differences, we explored whether the presence or absence of IA affects affective or cognitive abilities. Independent t-tests revealed no significant differences between these groups in terms of depression, anxiety, attention, verbal and non-verbal episodic memory, language, or working memory (t-test, p-values > 0.1).

### 3.5. Genetic influence on the IA

To investigate the potential genetic influence on IA development, its prevalence was compared across different genetic groups: monozygotic twins (MZ), dizygotic twins (DZ), non-twin siblings, and unrelated individuals.

After excluding “kissing” thalami cases, the sample included 175 MZ twins, 98 DZ twins, 146 non-twin siblings, and 117 unrelated individuals. The exclusions of subjects paired with those having a kissing thalamus were not applied to retain a larger effective sample size for subsequent sociodemographic analyses, which did not rely on paired data. Biological sex and education levels did not differ significantly between the genetic groups (χ² test = 3.10, p-value = 0.38; ANOVA, p-value > 0.05). However, non-twin siblings were slightly younger than MZ twins and unrelated subjects (27.1 vs. 29.3 years old; Dunn’s test with Bonferroni correction, p-value < 0.001). No significant variation in IA prevalence was observed between the groups (χ² test = 3.10, p-value = 0.38) (Table 1). Notably, the non-twin siblings group included 41 mixed-gender pairs, the unrelated group included 23 mixed-gender pairs, while no mixed-gender pairs were present in the MZ and DZ twin groups.

**Table 1:** Demographic data by genetic groups.

|  | MONOZYGOTIC | DIZYGOTIC | NOT TWIN<br>SIBLINGS | UNRELATED |
| --- | --- | --- | --- | --- |
| N | 175 | 98 | 146 | 117 |
| FEMALE | 57.0% | 54.0% | 47.3% | 54.7% |
| MEAN AGE ± SD | 29.3 ± 3.3 | 28.6 ± 3.4 | 27.1 ± 3.7 | 29.1 ± 3.7 |
| MEAN YEARS OF EDUCATION ± SD | 15.1 ± 1.8 | 15.3 ± 1.6 | 14.8 ± 1.9 | 14.8 ± 1.8 |
| IA PRESENCE (%) | 89.7 | 82.7 | 88.4 | 86.3 |

For the following analyses involving pairs of subjects, exclusions were made for related “kissing” cases (subjects paired with those having a kissing thalamus) and one unmatched unrelated subject, resulting in 164 monozygotic twins, 94 dizygotic twins, 136 non-twin siblings, and 116 unrelated subjects.

A logistic regression model was applied to determine whether each pair of subjects shared the same IA characterization by groups. The results revealed a significant difference between the genetic groups in predicting the presence or absence of IA between paired subjects (χ² test = 32, p-value < 0.001). The model’s accuracy in predicting the dependent variable was 86%. The odds ratios were 7 for the MZ group, 3.7 for the DZ group, 0.35 for the Non-Twin group, and 0.33 for the Unrelated group. This indicates a strong effect in the MZ group on the probability of having the same IA presence/absence, while no such effect was observed in the other groups. MZ twins had a 96% concordance rate for IA presence/absence, significantly higher than DZ twins (79%), Non-Twin siblings (85%), and Unrelated individuals (75%) (χ² test = 28, p-value < 0.001) (Fig. 4).

Interestingly, 100% of female MZ twins had the same IA characterization, while three pairs of male twins exhibited different IA characteristics (χ² test = 8.8, p-value = 0.004). Among these male pairs, those without an IA showed an average increase in third ventricle volume of 122 mm³.

### 3.6 Genetic influence on the IA variants

We then investigated whether genetic factors influenced specific IA anatomical variants. Due to their low frequency, statistical analysis was not conducted for the “Bilobar” and “Double” variants. No effects of age, years of education, or manual laterality were found between groups (Kruskal-Wallis test; p-value > 0.5). However, biological sex significantly influenced IA type, with a higher prevalence of broad IA in females (χ² test, p-value < 0.001, effect size = 17.4). No significant differences in the proportion of IA variants were observed between genetic groups (χ² test = 6.6, p-value > 0.5) (Tab. 2).

**Table 2:** Demographic data by IA anatomical variants.

|  | One | Broad | Double | Bilobar | Absent |
| --- | --- | --- | --- | --- | --- |
| All subjects | 328 (61%) | 132 (25%) | 4 (<1%) | 4 (<1%) | 68 (13%) |
| Mean age ± SD | 28.5 ± 3.7 | 28.6 ± 3.5 | 25.8 ± 3.9 | 30.3 ± 4.2 | 28.6 ± 3.3 |
| Mean years of education ± SD | 15 ± 1.7 | 14.9 ± 1.8 | 14.5 ± 1.3 | 16.3 ± 0.5 | 15 ± 1.7 |
| Female (%) | 51.2 | 72.0 | 50.0 | 75.0 | 26.5 |
| MZ (%) | 61.1 | 27.4 | <1 | <1 | 10.3 |
| DZ (%) | 56.1 | 24.5 | 1 | 1 | 17.4 |
| Not Twin (%) | 65.1 | 21.9 | 1.4 | 0 | 11.6 |
| Unrelated (%) | 60.7 | 23.9 | 0 | 1.7 | 13.7 |

Genetics’ effect on the adhesion type was evaluated using paired data between siblings and unrelated subjects through logistic regression. Results showed a significant difference between the genetic groups concerning the ability to predict having the same IA variants between paired subjects (χ² test = 21, p-value < 0.001). The accuracy of the model indicating the ability to predict having the same IA variant was 68%. The odds Ratio for the MZ group was 3.3, 1.3 in DZ, 1.4 for the Not Twin Group, and 1.2 for the Unrelated group. It indicates a slight effect of the group MZ on the probability of having the same IA, while no such effect was observed in other groups. Rates of pairs of subjects having the same IA in each group were 81% for MZ, 56.4% for DZ, 64% for Not Twin, and 60% for Unrelated, which highlights higher randomness among all groups except MZ. A Chi-squared test was conducted on the ability of each group to predict having the same IA variants and confirmed a significant difference (χ² test = 20.5, p-value < 0.001) (Fig.5). The MZ group had a significantly better prediction of having the same IA compared to the three other groups without differences between biological sex (χ² test = 0.002, p-value > 0.05).

**Figure 5:**
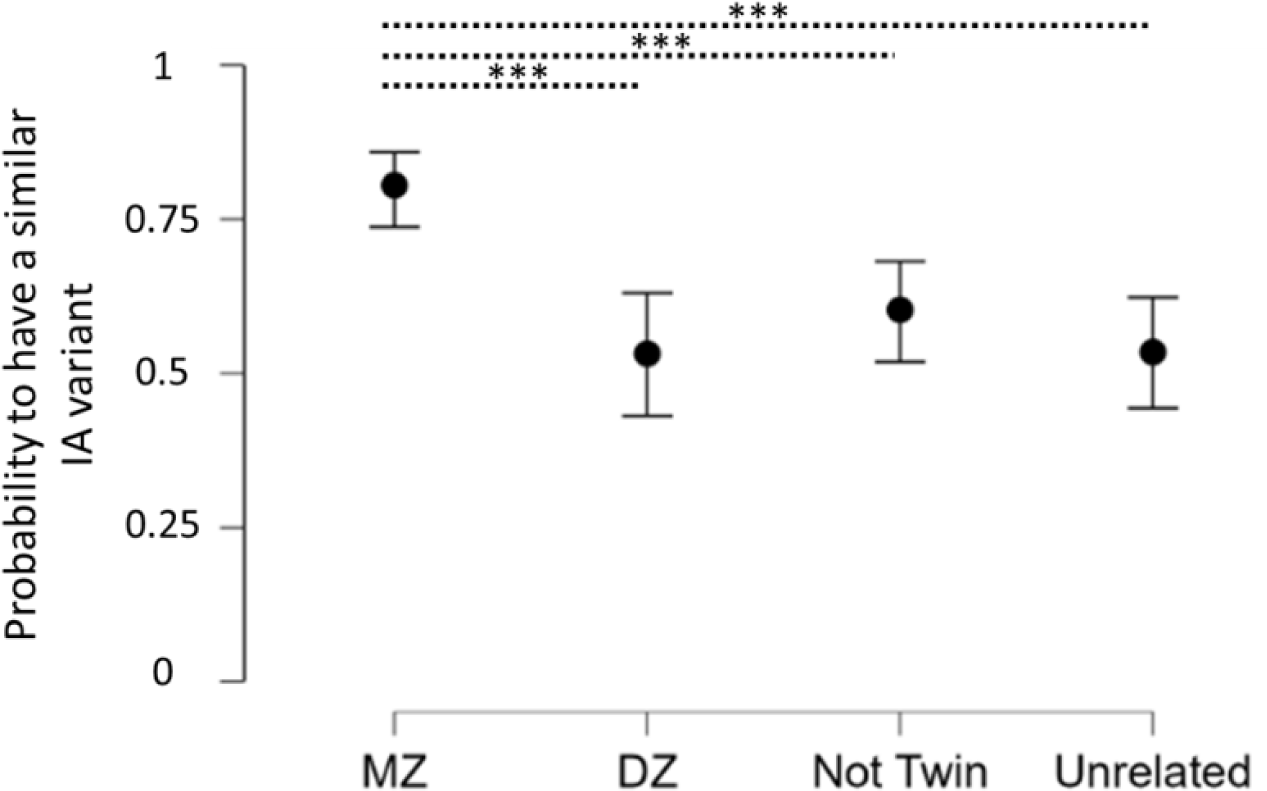
probability of having the same IA between each sibling or randomly paired unrelated subjects by genetic groups (MZ: monozygotic, DZ: dizygotic) ± 95% confidence interval. χ² p-values corrected for Bonferroni corrections: *** < 0.001, ** < 0.005.

## 4. Discussion

Leveraging a large cohort of 591 subjects, we found that the IA was more prevalent in females (94%) than males (80%). These differences were associated with a larger third ventricle, smaller corpus callosum, and cortical thinning in several regions in subjects without an IA. Despite these anatomical variations, no significant neuropsychological difference was evidenced. This study also reveals, for the first time, genetic influences on both the presence and anatomical variations of IA, with notable sexual dimorphism. Specifically, females exhibited a higher prevalence of IA, more broad anatomical variants, and there was 100% concordance in IA presence among female MZ twins, highlighting a significant biological sex impact on the IA.

### Prevalehnce of IA

Consistent with previous literature, the IA was present in about 87% of subjects, with a higher absence rate in males (20%) compared to females (6%) (Rabl 1958; Malobabic et al. 1987; Nopoulos et al. 2001; Guillery and Sherman 2002; Takahashi et al. 2008; Borghei et al. 2020, 2021; Wong et al. 2021; Sahin et al. 2023, Vidal et al. 2024). No significant differences were found in age, education, or manual laterality. While our dataset of 591 subjects is the largest in IA literature involving both neuroimaging and neuropsychological assessments in healthy individuals (e.g., Borghei et al. 2020: 402; Damle et al. 2021: 233), the age range of our participants is limited, spanning from 22 to 35 years. Despite this limitation, our findings align with Borghei et al. 2020, who found no age effects. In contrast, Damle et al. 2021, who included a broader age range of 8 to 68 years, could observe age-related influences on IA characteristics and cognitive performance. This highlights the need for future studies with a broader age range to understand these relationships better.

### Anatomical Differences

Structural analyses revealed significant differences between individuals with and without an IA. The absence of IA was associated with increased cerebrospinal fluid volumes and enlarged third ventricles in individuals without an IA. This finding aligns with earlier studies linking ventricle enlargement to the absence of IA, even more among patients with schizophrenia (Meisenzahl et al. 2002; Takahashi et al. 2010). For example, among the three pairs of male monozygotic twins with differing IA characteristics reported in our study, the sibling without an IA systematically had a larger third ventricle volume.

Additionally, individuals without IA exhibited reduced bi-thalamic and corpus callosum volumes alongside cortical thinning in several areas. These included the right entorhinal cortex, bilateral cuneus and precuneus, pericalcarine, insula, and lateral and medial orbitofrontal regions. This suggests that those regions are cortical areas of IA projections via both thalami, aligning with previous tractography studies (Damle et al. 2017; Kochanski et al. 2018; Borghei et al. 2020, 2021; Sahin et al. 2023). However, other structures were also found to be structurally connected with the IA in the literature, such as the amygdala, hippocampus, accumbens, and caudate nucleus, which we did not identify in this study (Damle et al. 2017; Kochanski et al. 2018; Borghei et al. 2020, 2021; Sahin et al. 2023). In addition to the literature and among the most significant differences in the absence of an IA, we also identified thinning in the bilateral parietal, precentral and paracentral regions, supramarginal, superior frontal cortices, and the banks of the superior temporal sulcus, pars opercularis, triangularis, and orbitalis. Connecting those findings with atlas-guided tract reconstruction and suggestions that thalamic radiations are the areas through which IA fibers reach cortical areas (Damle et al. 2017; Kochanski et al. 2018; Borghei et al. 2020, 2021), the pars opercularis, triangularis, and orbitalis are indeed known projection areas of the anterior thalamic radiations, while posterior thalamic radiations project to the parietal and occipital lobes. The superior thalamic radiations connect the thalamus to the precentral and paracentral gyri, and the inferior thalamic radiations to the insula, temporal, and frontal lobes (Zhang et al. 2010).

### Neuropsychological and affective impact of the presence or absence of the IA

Despite these anatomical differences, this study did not find significant neuropsychological or affective difference associated with IA presence/absence in this group of healthy individuals. The different studies that have tried to find neuropsychological differences between groups of healthy subjects with and without an IA have also usually failed to report any difference (Vidal et al. 2024), except marginally (Damle et al. 2017; Borghei et al. 2020). It is possible that the psychological tests used in the HCP and the other studies may not be sensitive enough to detect subtle differences in healthy subjects, all the more so as there is yet no known specific function associated with the IA that could be tested. The HCP group of subjects used in this project was also young (22-35 years old) and homogeneous, further suggesting that subjects might have performed at the ceiling. It is possible that using more sensitive tests in groups of older subjects could reveal subtler differences.

In addition, it is possible that compensatory neural mechanisms might mitigate potential cognitive impacts of IA presence/absence, supporting the hypothesis that the brain can reorganize itself to maintain cognitive function in the absence of IA (Vidal et al. 2024). IA functions could also be taken over by other commissural pathways, such as the corpus callosum or the anterior or posterior commissures, in case of IA absence (Borghei et al. 2021; Kochanski et al. 2017). Overall, associating the IA with specific cognitive functions might prove difficult.

### Genetic Influence on IA Prevalence and Anatomical Variants

The results of this study highlight a significant genetic influence on the presence and anatomical variations of the IA. Monozygotic (MZ) twins exhibited a higher concordance rate of IA presence (96%), compared to dizygotic (DZ) twins (79%), non-twin siblings (85%), or unrelated subjects (75%). However, 4% of subjects in the MZ group did not share the same IA characterization as their siblings, potentially implicating biological sex effects since these discrepancies were only found among men.

This analysis aligns with previous studies that highlight strong anatomical correlations between MZ twins compared to matched controls, underscoring the genetic contribution to both prenatal and postnatal brain development (White et al. 2002). However, MZ twins can exhibit differential diagnoses of neurodevelopmental disorders such as schizophrenia, bipolar disorder, and attention deficit hyperactivity disorder (ADHD). Notably, in schizophrenia, the affected twin more often shows third ventricle enlargement, indicating anatomical differences not solely explained by genetics (Suddath et al. 1990). Among discordant twins for these conditions, various anatomical differences have been observed alongside epigenetic variations (Chen et al. 2008; Dempster et al. 2011). For instance, differential X-chromosome inactivation has been suggested in MZ female twins discordant for bipolar disorder (Rosa et al. 2008), and epigenetic changes, such as alterations in DNA methylation of the dopamine D2 receptor gene or the catechol-O-methyltransferase gene, have been noted in MZ twins discordant for schizophrenia (Petronis et al. 2004).

### Biological Sex Differences and Genetic Factors Modulating IA Prevalence

Biological sex emerged as a significant factor of IA presence, with females exhibiting a higher prevalence of IA than males, in robust agreement with previous studies (Rabl 1958; Malobabic et al. 1987; Erbagci et al. 2002; Guillery and Sherman 2002; Damle et al. 2017; Borghei et al. 2020; Tsutsumi et al. 2021; Sahin et al. 2023; Vidal et al. 2024). Consistent with more recent findings, a higher rate of broad IA in females was also demonstrated in this study (Trzesniak et al. 2016; Pavlovic et al. 2020; Tsutsumi et al. 2021; Patra et al. 2022).

In line with these particular features present only in females, our novel finding shows 100% concordance in IA presence among female MZ twins (compared to 96% for males). It suggests a genetic predisposition for IA that is either carried by autosomes or by the X chromosome, as both biological sexes share it. However, sexual dimorphism likely influences IA presence.

Although males’ and females’ brains are very similar, differences exist. On average, some studies indicate that male brains weigh more than female brains, resulting in proportionally larger cerebrospinal fluid, intracranial volume, gray matter, white matter, and ventricular volumes (Ruigrok et al. 2014). There is a great disparity in the literature concerning cortical or subcortical volumes, cortical thicknesses, and brain lateralization dimorphisms. A recent meta-synthesis demonstrated no reliable subcortical volumes or cortical thickness differences when adjusting for brain size, nor lateralization dimorphisms (Eliot et al. 2021) despite more uncertainty about slightly larger amygdala, hippocampus, insula, or putamen among males (Ruigrok et al. 2014).

Cortical volume studies lack consistency, but recent findings with corrected volumes show larger male volume in the anterior parahippocampal cortex and larger female volume in the prefrontal, inferior parietal, superior temporal, and cingulate cortices (Ruigrok et al. 2014; Potvin et al. 2018; Eliot et al. 2021). Structural connection studies also show discrepancies, but females tend to have a slightly larger corpus callosum and anterior commissure (Choi et al. 2011; Potvin et al. 2016, 2018). These larger structures in females are demonstrated in the present study to be larger in individuals with IA, suggesting a need to refine further research to understand these differences. Currently, the only consistently reliable difference between males and females is the different IA prevalence and anatomical variant (Eliot et al. 2021).

To consider this IA and anatomical variants prevalence sex dimorphism, it has been demonstrated that sex-chromosomal (sex-specific genes on the sex chromosome) and sex-hormone effects (estrogen, androgens, etc.) play important roles during brain development. Any imbalance could generate sex-related anatomical and functional differences, leading to different vulnerabilities to behavioral, neurodevelopmental, and neurodegenerative diseases (Gilies and McArthur 2010). Males are almost twice as likely to develop neurodevelopmental diseases such as autism spectrum disorders, attention deficit hyperactivity disorder, schizophrenia, and dyslexia or neurodegenerative diseases such as Parkinson’s disease (Wooten et al. 2004; Gillberg et al. 2006; Balint et al. 2009; Boyle et al. 2011). Some of those disorders etiologically involve the midbrain dopaminergic system (Aleman et al. 2003; Hennessy et al. 2004; Barnes et al. 2005; Baron-Cohen et al. 2005; Shulman and Bhat 2006; Becker and Hu 2008). Among pediatric patients with midline abnormalities, IA absence is four times higher compared to healthy groups (Whitehead and Najim 2020) and twice higher in schizophrenia spectrum disorders. Preceding these results, Tresniak et al. 2011 suggested an IA role of the dopaminergic system in schizophrenia spectrum disorders etiopathology (Andreasen et al. 1994, Buchsbaum et al. 2006; Ettinger et al. 2013).

Bridging the link between IA, the dopaminergic system, and sexual dimorphism, three hypotheses can be stated using sex hormones, sex chromosomes, and/or epigenetic dimorphism effects. The first hypothesis could leverage the sex-hormone effects theory. In the absence of estrogen synthesis, apoptosis of dopaminergic neurons occurs spontaneously in the adult male mouse hypothalamus (Hill et al. 2004). Estrogens also have neuroprotective effects in Parkinson’s disease via the nigrostriatal dopaminergic system and improve schizophrenic symptoms and therapeutic responses to treatments when increased (Quinn and Marsden 1986; Kulkani et al. 2001; Baron-Cohen et al. 2005; Czech et al. 2014). Males might be more vulnerable to the IA loss during development and lifespan due to the protective and/or pro-dopaminergic effects of estrogen. For the sexual chromosomal theory, SRY (Sex-determining Region on the Y chromosome) is an interesting candidate as it is expressed in dopamine-abundant brain regions, regulating dopamine biosynthesis in males. Its dysregulation may causally induce Parkinson’s disease (Czech et al. 2014). SRY dysregulation among males during early development could alter the dopaminergic system, leading to IA anomalies. Finally, epigenetic studies indicate sex-specific signals induced by both environmental and preprogrammed hormonal or genetic cues (Gatewood et al. 2006; Barker et al. 2010; Cox and Rissman 2011; McCarthy and al. 2012), causing differential gene expression between males and females during development and across the lifespan (McCarthy and al. 2012, Yang and Shah 2014). These epigenetic phenomena can impact the brain even after the perinatal window closes (Nugent and Bale 2015).

### Significance

Overall, our results support the hypothesis that IA presence possesses a genetic predisposition. IA absence, rather than having a direct causal pathological effect, could be an indicator or a marker of abnormal neurodevelopment of midline structures involving mechanisms for which males are more vulnerable. Associated with other abnormalities, IA absence could possibly be related to structural and functional aberrations such as deficits in the dopaminergic system and could involve a higher risk of developing a neuropsychiatric disorder (Nopoulos et al. 2001; Trzesniak et al. 2011). Being in possession of an IA would then indicate the integrity of adjacent midline structures and the third ventricle, reflecting a greater probability of better outcomes. As such, identifying the presence or absence of the IA in neurodevelopmental diseases appears highly promising, as it is generally easy to assess on good-quality MRI and could be a useful biomarker.

### Limits

This article presents several limitations. The association between IA and age could not be identified due to the limited age distribution of subjects. Additionally, the low number of siblings with different IA characterizations in the MZ group prevented proper neuropsychological statistical analyses. Finally, the DZ group did not include mixed biological sex sibling pairs for reasons remaining unclear to us. The Not-Twin group was the only one with mixed-biological sex sibling pairs.

### Conclusion

This study provides robust evidence that genetic factors significantly influence IA prevalence and anatomical variants. While IA absence correlates with specific neuroanatomical differences, it does not appear to impact cognitive function in healthy individuals. Future research should focus on the clinical significance of IA absence, its potential role as a biomarker for neurodevelopmental abnormalities, as well as the possible mechanisms underlying compensatory neural pathways that preserve cognitive function in the absence of IA.

## 5. Acknowledgments

We want to express our gratitude to Vinod Kumar for his assistance in obtaining access to the HCP database.

## 6. Authors contributions

J.P.V. and E.J.B. designed the study.

J.P.V. performed the computations and analysis under the supervision of L.D. and E.J.B. and wrote the manuscript with their support.

J.P, P.P, J.P., L.D, E.J.B substantively revised the manuscript.

## 8. Data availability

All data are provided by the Human Connectome Project (HCP) and are either part of a public dataset or shared on request.

## Supplementary data

**Supplementary Table 1:**
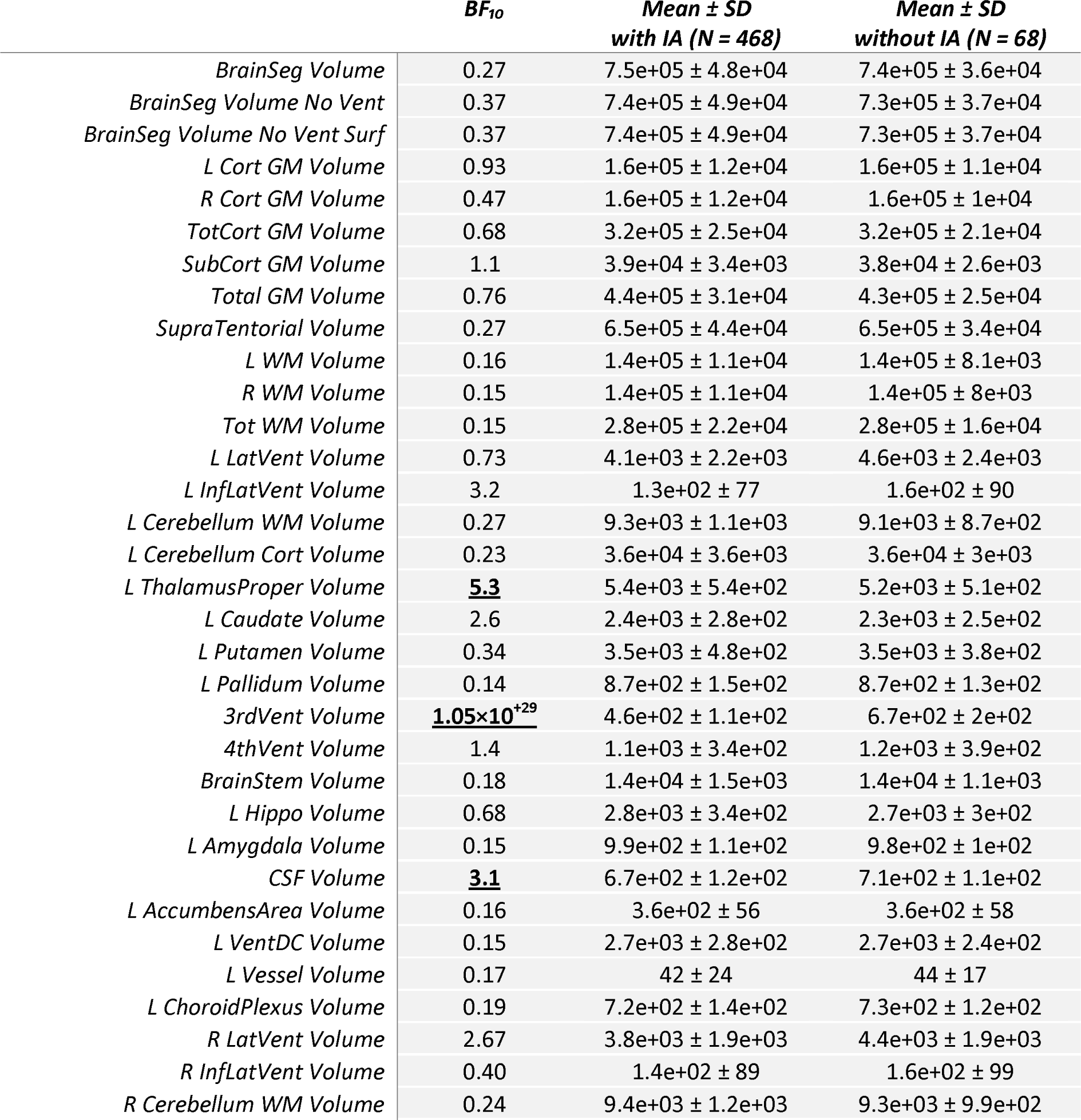
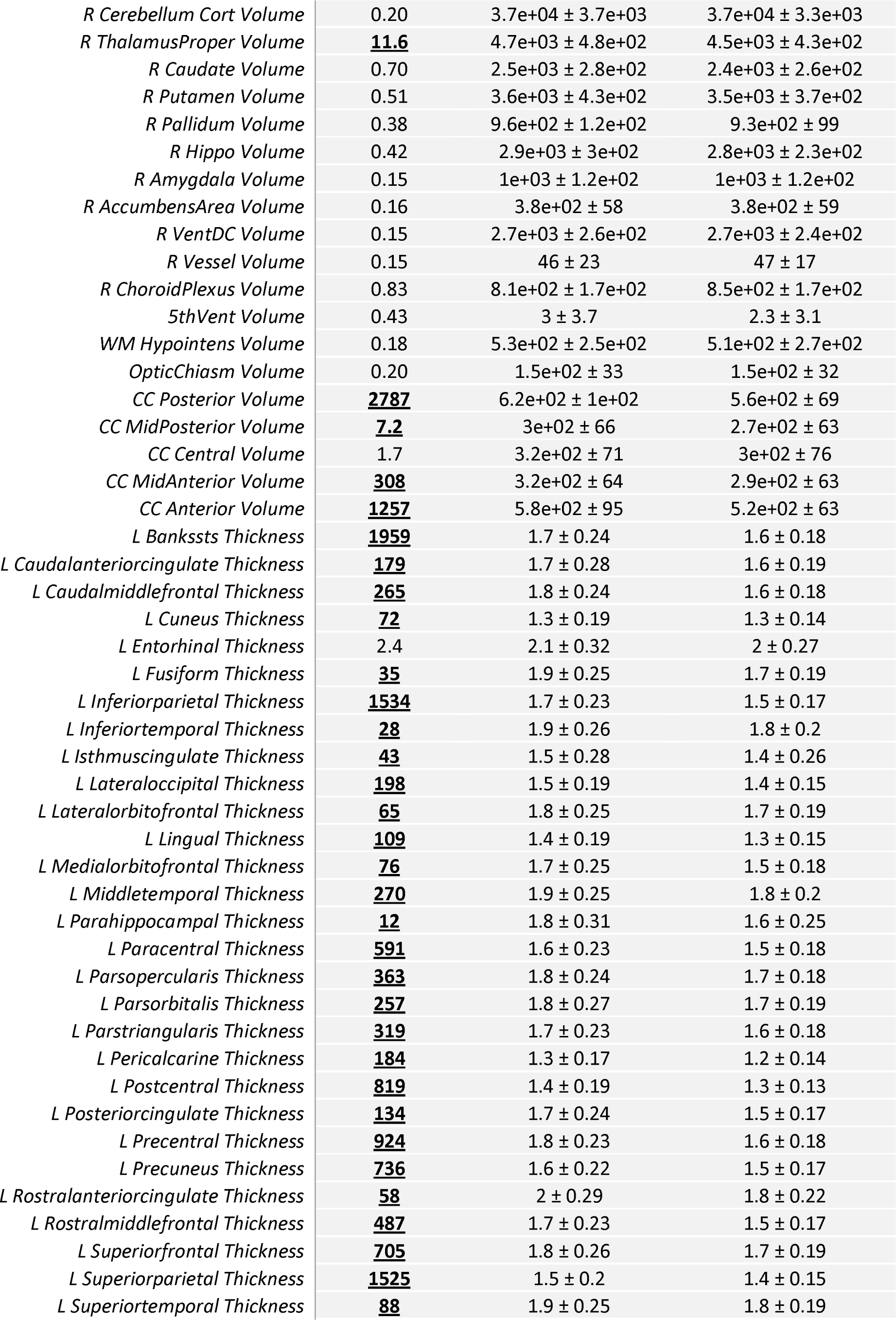
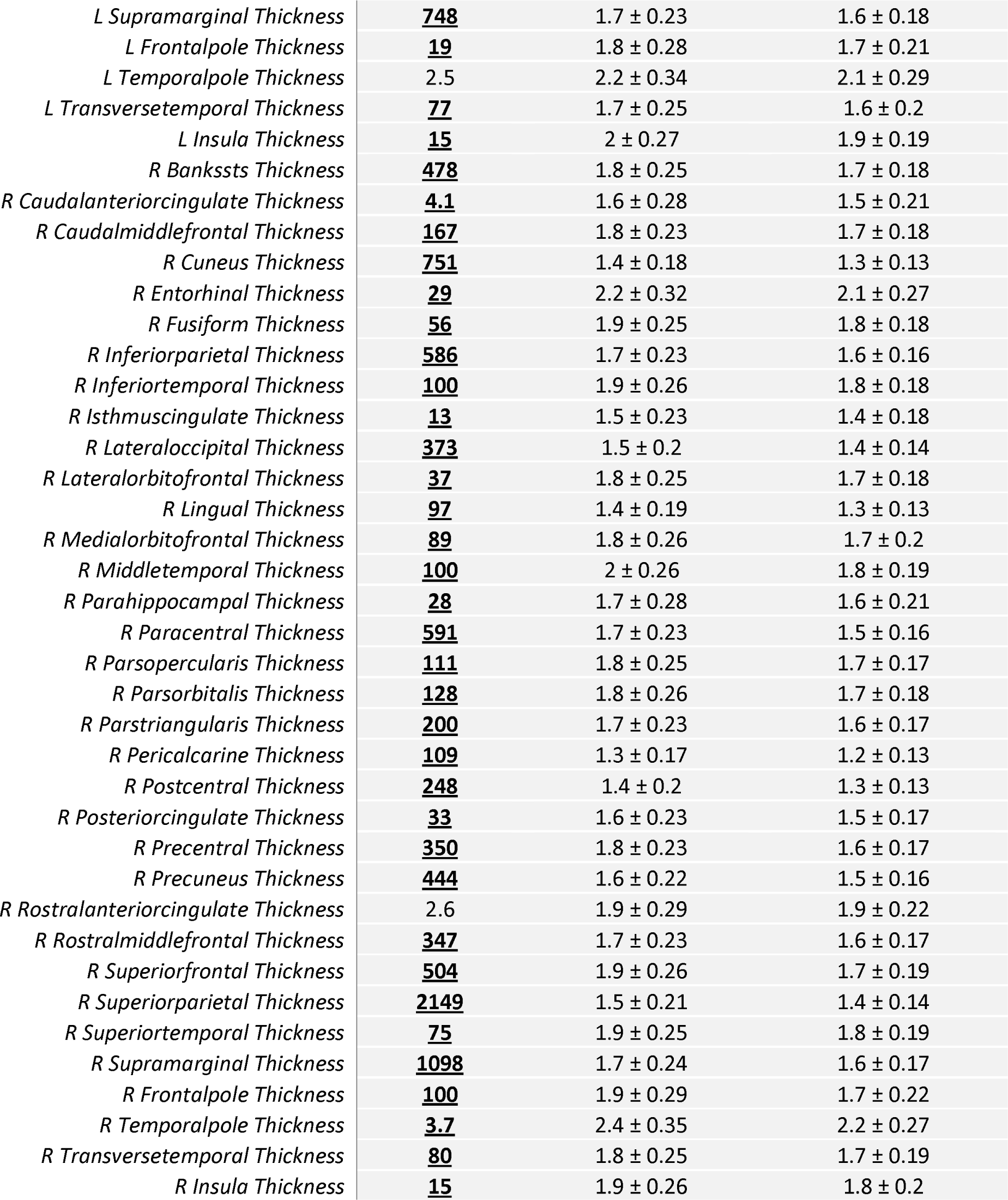
Mean ± SD of volumes and cortical thicknesses computed using Freesurfer software for all subjects with (n = 468) or without an IA (n = 68). Freesurfer nomenclature is maintained for consistency. Bayes Factor (BF_10_) resulting from a Bayesian t-test are included. A BF_10_ greater than 3 indicates at least moderate evidence of a difference between the two groups and is represented in bold and underlined.

|  | **BF<sub>10</sub>** | **Mean ± SD**<br>**with IA (N = 468)** | **Mean ± SD**<br>**without IA (N = 68)** |
| --- | --- | --- | --- |
| *BrainSeg Volume* | 0.27 | 7.5e+05 ± 4.8e+04 | 7.4e+05 ± 3.6e+04 |
| *BrainSeg Volume No Vent* | 0.37 | 7.4e+05 ± 4.9e+04 | 7.3e+05 ± 3.7e+04 |
| *BrainSeg Volume No Vent Surf* | 0.37 | 7.4e+05 ± 4.9e+04 | 7.3e+05 ± 3.7e+04 |
| *L Cort GM Volume* | 0.93 | 1.6e+05 ± 1.2e+04 | 1.6e+05 ± 1.1e+04 |
| *R Cort GM Volume* | 0.47 | 1.6e+05 ± 1.2e+04 | 1.6e+05 ± 1e+04 |
| *TotCort GM Volume* | 0.68 | 3.2e+05 ± 2.5e+04 | 3.2e+05 ± 2.1e+04 |
| *SubCort GM Volume* | 1.1 | 3.9e+04 ± 3.4e+03 | 3.8e+04 ± 2.6e+03 |
| *Total GM Volume* | 0.76 | 4.4e+05 ± 3.1e+04 | 4.3e+05 ± 2.5e+04 |
| *SupraTentorial Volume* | 0.27 | 6.5e+05 ± 4.4e+04 | 6.5e+05 ± 3.4e+04 |
| *L WM Volume* | 0.16 | 1.4e+05 ± 1.1e+04 | 1.4e+05 ± 8.1e+03 |
| *R WM Volume* | 0.15 | 1.4e+05 ± 1.1e+04 | 1.4e+05 ± 8e+03 |
| *Tot WM Volume* | 0.15 | 2.8e+05 ± 2.2e+04 | 2.8e+05 ± 1.6e+04 |
| *L LatVent Volume* | 0.73 | 4.1e+03 ± 2.2e+03 | 4.6e+03 ± 2.4e+03 |
| *L InfLatVent Volume* | 3.2 | 1.3e+02 ± 77 | 1.6e+02 ± 90 |
| *L Cerebellum WM Volume* | 0.27 | 9.3e+03 ± 1.1e+03 | 9.1e+03 ± 8.7e+02 |
| *L Cerebellum Cort Volume* | 0.23 | 3.6e+04 ± 3.6e+03 | 3.6e+04 ± 3e+03 |
| *L ThalamusProper Volume* | **<u>5.3</u>** | 5.4e+03 ± 5.4e+02 | 5.2e+03 ± 5.1e+02 |
| *L Caudate Volume* | 2.6 | 2.4e+03 ± 2.8e+02 | 2.3e+03 ± 2.5e+02 |
| *L Putamen Volume* | 0.34 | 3.5e+03 ± 4.8e+02 | 3.5e+03 ± 3.8e+02 |
| *L Pallidum Volume* | 0.14 | 8.7e+02 ± 1.5e+02 | 8.7e+02 ± 1.3e+02 |
| *3rdVent Volume* | **<u>1.05×10<sup>+29</sup></u>** | 4.6e+02 ± 1.1e+02 | 6.7e+02 ± 2e+02 |
| *4thVent Volume* | 1.4 | 1.1e+03 ± 3.4e+02 | 1.2e+03 ± 3.9e+02 |
| *BrainStem Volume* | 0.18 | 1.4e+04 ± 1.5e+03 | 1.4e+04 ± 1.1e+03 |
| *L Hippo Volume* | 0.68 | 2.8e+03 ± 3.4e+02 | 2.7e+03 ± 3e+02 |
| *L Amygdala Volume* | 0.15 | 9.9e+02 ± 1.1e+02 | 9.8e+02 ± 1e+02 |
| *CSF Volume* | **<u>3.1</u>** | 6.7e+02 ± 1.2e+02 | 7.1e+02 ± 1.1e+02 |
| *L AccumbensArea Volume* | 0.16 | 3.6e+02 ± 56 | 3.6e+02 ± 58 |
| *L VentDC Volume* | 0.15 | 2.7e+03 ± 2.8e+02 | 2.7e+03 ± 2.4e+02 |
| *L Vessel Volume* | 0.17 | 42 ± 24 | 44 ± 17 |
| *L ChoroidPlexus Volume* | 0.19 | 7.2e+02 ± 1.4e+02 | 7.3e+02 ± 1.2e+02 |
| *R LatVent Volume* | 2.67 | 3.8e+03 ± 1.9e+03 | 4.4e+03 ± 1.9e+03 |
| *R InfLatVent Volume* | 0.40 | 1.4e+02 ± 89 | 1.6e+02 ± 99 |
| *R Cerebellum WM Volume* | 0.24 | 9.4e+03 ± 1.2e+03 | 9.3e+03 ± 9.9e+02 |
| *R Cerebellum Cort Volume* | 0.20 | 3.7e+04 ± 3.7e+03 | 3.7e+04 ± 3.3e+03 |
| *R ThalamusProper Volume* | **<u>11.6</u>** | 4.7e+03 ± 4.8e+02 | 4.5e+03 ± 4.3e+02 |
| *R Caudate Volume* | 0.70 | 2.5e+03 ± 2.8e+02 | 2.4e+03 ± 2.6e+02 |
| *R Putamen Volume* | 0.51 | 3.6e+03 ± 4.3e+02 | 3.5e+03 ± 3.7e+02 |
| *R Pallidum Volume* | 0.38 | 9.6e+02 ± 1.2e+02 | 9.3e+02 ± 99 |
| *R Hippo Volume* | 0.42 | 2.9e+03 ± 3e+02 | 2.8e+03 ± 2.3e+02 |
| *R Amygdala Volume* | 0.15 | 1e+03 ± 1.2e+02 | 1e+03 ± 1.2e+02 |
| *R AccumbensArea Volume* | 0.16 | 3.8e+02 ± 58 | 3.8e+02 ± 59 |
| *R VentDC Volume* | 0.15 | 2.7e+03 ± 2.6e+02 | 2.7e+03 ± 2.4e+02 |
| *R Vessel Volume* | 0.15 | 46 ± 23 | 47 ± 17 |
| *R ChoroidPlexus Volume* | 0.83 | 8.1e+02 ± 1.7e+02 | 8.5e+02 ± 1.7e+02 |
| *5thVent Volume* | 0.43 | 3 ± 3.7 | 2.3 ± 3.1 |
| *WM Hypointens Volume* | 0.18 | 5.3e+02 ± 2.5e+02 | 5.1e+02 ± 2.7e+02 |
| *OpticChiasm Volume* | 0.20 | 1.5e+02 ± 33 | 1.5e+02 ± 32 |
| *CC Posterior Volume* | **<u>2787</u>** | 6.2e+02 ± 1e+02 | 5.6e+02 ± 69 |
| *CC MidPosterior Volume* | **<u>7.2</u>** | 3e+02 ± 66 | 2.7e+02 ± 63 |
| *CC Central Volume* | 1.7 | 3.2e+02 ± 71 | 3e+02 ± 76 |
| *CC MidAnterior Volume* | **<u>308</u>** | 3.2e+02 ± 64 | 2.9e+02 ± 63 |
| *CC Anterior Volume* | **<u>1257</u>** | 5.8e+02 ± 95 | 5.2e+02 ± 63 |
| *L Bankssts Thickness* | **<u>1959</u>** | 1.7 ± 0.24 | 1.6 ± 0.18 |
| *L Caudalanteriorcingulate Thickness* | **<u>179</u>** | 1.7 ± 0.28 | 1.6 ± 0.19 |
| *L Caudalmiddlefrontal Thickness* | **<u>265</u>** | 1.8 ± 0.24 | 1.6 ± 0.18 |
| *L Cuneus Thickness* | **<u>72</u>** | 1.3 ± 0.19 | 1.3 ± 0.14 |
| *L Entorhinal Thickness* | 2.4 | 2.1 ± 0.32 | 2 ± 0.27 |
| *L Fusiform Thickness* | **<u>35</u>** | 1.9 ± 0.25 | 1.7 ± 0.19 |
| *L Inferiorparietal Thickness* | **<u>1534</u>** | 1.7 ± 0.23 | 1.5 ± 0.17 |
| *L Inferiortemporal Thickness* | **<u>28</u>** | 1.9 ± 0.26 | 1.8 ± 0.2 |
| *L Isthmuscingulate Thickness* | **<u>43</u>** | 1.5 ± 0.28 | 1.4 ± 0.26 |
| *L Lateraloccipital Thickness* | **<u>198</u>** | 1.5 ± 0.19 | 1.4 ± 0.15 |
| *L Lateralorbitofrontal Thickness* | **<u>65</u>** | 1.8 ± 0.25 | 1.7 ± 0.19 |
| *L Lingual Thickness* | **<u>109</u>** | 1.4 ± 0.19 | 1.3 ± 0.15 |
| *L Medialorbitofrontal Thickness* | **<u>76</u>** | 1.7 ± 0.25 | 1.5 ± 0.18 |
| *L Middletemporal Thickness* | **<u>270</u>** | 1.9 ± 0.25 | 1.8 ± 0.2 |
| *L Parahippocampal Thickness* | **<u>12</u>** | 1.8 ± 0.31 | 1.6 ± 0.25 |
| *L Paracentral Thickness* | **<u>591</u>** | 1.6 ± 0.23 | 1.5 ± 0.18 |
| *L Parsopercularis Thickness* | **<u>363</u>** | 1.8 ± 0.24 | 1.7 ± 0.18 |
| *L Parsorbitalis Thickness* | **<u>257</u>** | 1.8 ± 0.27 | 1.7 ± 0.19 |
| *L Parstriangularis Thickness* | **<u>319</u>** | 1.7 ± 0.23 | 1.6 ± 0.18 |
| *L Pericalcarine Thickness* | **<u>184</u>** | 1.3 ± 0.17 | 1.2 ± 0.14 |
| *L Postcentral Thickness* | **<u>819</u>** | 1.4 ± 0.19 | 1.3 ± 0.13 |
| *L Posteriorcingulate Thickness* | **<u>134</u>** | 1.7 ± 0.24 | 1.5 ± 0.17 |
| *L Precentral Thickness* | **<u>924</u>** | 1.8 ± 0.23 | 1.6 ± 0.18 |
| *L Precuneus Thickness* | **<u>736</u>** | 1.6 ± 0.22 | 1.5 ± 0.17 |
| *L Rostralanteriorcingulate Thickness* | **<u>58</u>** | 2 ± 0.29 | 1.8 ± 0.22 |
| *L Rostralmiddlefrontal Thickness* | **<u>487</u>** | 1.7 ± 0.23 | 1.5 ± 0.17 |
| *L Superiorfrontal Thickness* | **<u>705</u>** | 1.8 ± 0.26 | 1.7 ± 0.19 |
| *L Superiorparietal Thickness* | **<u>1525</u>** | 1.5 ± 0.2 | 1.4 ± 0.15 |
| *L Superiortemporal Thickness* | **<u>88</u>** | 1.9 ± 0.25 | 1.8 ± 0.19 |
| *L Supramarginal Thickness* | **<u>748</u>** | 1.7 ± 0.23 | 1.6 ± 0.18 |
| *L Frontalpole Thickness* | **<u>19</u>** | 1.8 ± 0.28 | 1.7 ± 0.21 |
| *L Temporalpole Thickness* | **2.5** | 2.2 ± 0.34 | 2.1 ± 0.29 |
| *L Transversetemporal Thickness* | **<u>77</u>** | 1.7 ± 0.25 | 1.6 ± 0.2 |
| *L Insula Thickness* | **<u>15</u>** | 2 ± 0.27 | 1.9 ± 0.19 |
| *R Bankssts Thickness* | **<u>478</u>** | 1.8 ± 0.25 | 1.7 ± 0.18 |
| *R Caudalanteriorcingulate Thickness* | **<u>4.1</u>** | 1.6 ± 0.28 | 1.5 ± 0.21 |
| *R Caudalmiddlefrontal Thickness* | **<u>167</u>** | 1.8 ± 0.23 | 1.7 ± 0.18 |
| *R Cuneus Thickness* | **<u>751</u>** | 1.4 ± 0.18 | 1.3 ± 0.13 |
| *R Entorhinal Thickness* | **<u>29</u>** | 2.2 ± 0.32 | 2.1 ± 0.27 |
| *R Fusiform Thickness* | **<u>56</u>** | 1.9 ± 0.25 | 1.8 ± 0.18 |
| *R Inferiorparietal Thickness* | **<u>586</u>** | 1.7 ± 0.23 | 1.6 ± 0.16 |
| *R Inferiortemporal Thickness* | **<u>100</u>** | 1.9 ± 0.26 | 1.8 ± 0.18 |
| *R Isthmuscingulate Thickness* | **<u>13</u>** | 1.5 ± 0.23 | 1.4 ± 0.18 |
| *R Lateraloccipital Thickness* | **<u>373</u>** | 1.5 ± 0.2 | 1.4 ± 0.14 |
| *R Lateralorbitofrontal Thickness* | **<u>37</u>** | 1.8 ± 0.25 | 1.7 ± 0.18 |
| *R Lingual Thickness* | **<u>97</u>** | 1.4 ± 0.19 | 1.3 ± 0.13 |
| *R Medialorbitofrontal Thickness* | **<u>89</u>** | 1.8 ± 0.26 | 1.7 ± 0.2 |
| *R Middletemporal Thickness* | **<u>100</u>** | 2 ± 0.26 | 1.8 ± 0.19 |
| *R Parahippocampal Thickness* | **<u>28</u>** | 1.7 ± 0.28 | 1.6 ± 0.21 |
| *R Paracentral Thickness* | **<u>591</u>** | 1.7 ± 0.23 | 1.5 ± 0.16 |
| *R Parsopercularis Thickness* | **<u>111</u>** | 1.8 ± 0.25 | 1.7 ± 0.17 |
| *R Parsorbitalis Thickness* | **<u>128</u>** | 1.8 ± 0.26 | 1.7 ± 0.18 |
| *R Parstriangularis Thickness* | **<u>200</u>** | 1.7 ± 0.23 | 1.6 ± 0.17 |
| *R Pericalcarine Thickness* | **<u>109</u>** | 1.3 ± 0.17 | 1.2 ± 0.13 |
| *R Postcentral Thickness* | **<u>248</u>** | 1.4 ± 0.2 | 1.3 ± 0.13 |
| *R Posteriorcingulate Thickness* | **<u>33</u>** | 1.6 ± 0.23 | 1.5 ± 0.17 |
| *R Precentral Thickness* | **<u>350</u>** | 1.8 ± 0.23 | 1.6 ± 0.17 |
| *R Precuneus Thickness* | **<u>444</u>** | 1.6 ± 0.22 | 1.5 ± 0.16 |
| *R Rostralanteriorcingulate Thickness* | **2.6** | 1.9 ± 0.29 | 1.9 ± 0.22 |
| *R Rostralmiddlefrontal Thickness* | **<u>347</u>** | 1.7 ± 0.23 | 1.6 ± 0.17 |
| *R Superiorfrontal Thickness* | **<u>504</u>** | 1.9 ± 0.26 | 1.7 ± 0.19 |
| *R Superiorparietal Thickness* | **<u>2149</u>** | 1.5 ± 0.21 | 1.4 ± 0.14 |
| *R Superiortemporal Thickness* | **<u>75</u>** | 1.9 ± 0.25 | 1.8 ± 0.19 |
| *R Supramarginal Thickness* | **<u>1098</u>** | 1.7 ± 0.24 | 1.6 ± 0.17 |
| *R Frontalpole Thickness* | **<u>100</u>** | 1.9 ± 0.29 | 1.7 ± 0.22 |
| *R Temporalpole Thickness* | **<u>3.7</u>** | 2.4 ± 0.35 | 2.2 ± 0.27 |
| *R Transversetemporal Thickness* | **<u>80</u>** | 1.8 ± 0.25 | 1.7 ± 0.19 |
| *R Insula Thickness* | **<u>15</u>** | 1.9 ± 0.26 | 1.8 ± 0.2 |

